# Target Gene Notebook: Connecting genetics and drug discovery

**DOI:** 10.1101/757690

**Authors:** Mary Pat Reeve, Andrew Kirby, Jamey Wierzbowski, Mark Daly, Janna Hutz

## Abstract

Target Gene Notebook was developed to enable more efficient linking of genetic associations to functional biological information. This process is essential to translating genetic insights into therapeutic hypotheses and, eventually, drug discovery. Although many public databases provide access to unfiltered genome annotations and genetic results, there was no existing tool to maintain group curation and integration with proprietary experimental data. We provide Target Gene Notebook freely via the MIT open-source license for the purposes of assisting therapeutic target evaluation and the creation of durable institutional and public knowledge bases. Implemented as a Java backend serving mainly Javascript content derived from gene-specific SQLite databases, Target Gene Notebook enables automated access to the most widely used sources of genetic association, expression and protein QTL data, provides intuitive interfaces to credible set and colocalization information, and enables comprehensive literature review and annotation by multiple users simultaneously to create a consistent target knowledgebase within an organization or across a consortium. TargetGeneNotebook is freely available from GitHub https://github.com/targetgenenotebook/tgn.git under the MIT open-source end-user license agreement and a live version of the interface is provided at http://tgn.broadinstitute.org/.

## Introduction

Human genetic research has reached an exciting time in which genetic and medical data from human populations can be used for validating drug targets as safe and effective, discovering biomarkers and surrogate endpoints, and stratifying patients in trials (Plenge 2013, Nelson 2015). Genomewide association study (GWAS) scans across diverse diseases and traits produce robust evidence of variants’ causal impact on disease. However, additional data must be integrated to understand the molecular mechanisms by which a given variant might impact a gene’s biological activity in order to establish a therapeutic hypothesis. While genetics therefore provides a promising path to identify safe, effective and relevant targets the data sources upon which such inferences can be built are diverse, disorganized and disconnected. Data from resources focusing on protein biochemistry, gene expression, model organisms, and pathways can offer clues to the functional relevance of new genetic insights. These data can often be integrated bioinformatically with annotation pipelines, but expert human assessment is still required to determine relevance to the therapeutic area of interest, to integrate knowledge that is not amenable to bioinformatic integration, and to resolve any conflicting data. Within interdisciplinary teams, capturing the curator’s assessments alongside the supporting data is key to disseminating knowledge uniformly across a team. However, while much attention has been focused on analytics within many of these individual functional domains, there is no existing publicly available tool which facilitates the acquisition and interpretation of these data together, or even ensures that all relevant sources have been reviewed.

Target Gene Notebook (TGN) was developed to fill the void of providing a durable knowledge base about a potential therapeutic target across a team. TGN provides three parallel supports to bring clarity on the information about a target:

- Computational tools to show the architecture of the region, including novel tracks of data showing genotype-phenotype associations, expression and protein QTLs (eQTLS and pQTLs) and visualization of linkage-disequilibrium (LD) blocks
- Curatorial tools which allow multiple users from different technical backgrounds to record and integrate their assessments of the data from publications, public databases or internal, proprietary sources
- Modular, open framework to enable sharing (within institutions, between institutions or publicly), to readily evolve with new data types and to incorporate and provide modules from other developers
- The combined computational and curatorial tools contained in TGN enable researchers to maximize the data for drug design and elucidation of biological mechanisms

## Design and Implementation

### Automated data collection from public repositories

Searching for and manually annotating information from the numerous sources providing genetic and functional information on targets is not only time consuming, but also potentially unending, as these sources are constantly being updated. To save the user time gathering this data, a database build script automates the gathering of available information on the target, taking advantage of the extensive data-source harmonization/integration provided by Ensembl (Zerbino, 2018). The current release of the integrated database build script follows this workflow:

For any arbitrary target-gene symbol, genomic data are harvested via the Ensembl API and REST servers:

1. The identity and exon structure of the target-gene’s canonical transcript are retrieved.
2. The relevant genomic interval is retrieved, typically as the footprint of the target-gene’s canonical transcript +/- a user-defined extension (default is set to +/- 50kb).
3. The canonical transcript and exon structure of other genes overlapping the genomic interval are retrieved.
4. Variation is retrieved across the genomic interval. To supplement this, all pairs of markers from the 1000 Genomes Phase 3 release (1KGp3) with r2 >= 0.1 within 500kb of each other are collected for each major 1KGp3 population (EUR, AFR, EAS, SAS, AMR), re-taining those LD pairs containing at least one of the directly-collected variants with additional variant and frequency information available via hyperlink to gnomAD (http://gnomad.broadinstitute.org). (Karczewski 2019)
5. Population-specific allele frequencies and canonical-transcript impacts are collected for all known variants.
6. Phenotype associations across the genomic interval from the NHGRI-EBI GWAS catalog are retrieved. Publicly available information for each corresponding PubMed ID is also recovered separately from NCBI.
7. Cis-acting eQTL results are recovered for all genes overlapping the defined genomic interval. Results are filtered for those passing tis-sue-specific significance thresholds (separately downloaded from GTEx (https://gtexportal.org, release V6p)), being within the defined genomic fragment, and being typed in 1KGp3 EUR.
8. Gene-phenotype associations from MIM morbid and Cancer Gene Census assigned to the defined genomic interval are recovered.
9. Allele-phenotype associations from OMIM and ClinVar that are assigned across the defined genomic fragment are recovered.
10. Tissue-specific expression-level graphics, based on GTEx data (separately downloaded from https://gtexportal.org, release V6p), are prepared for each gene overlapping the defined genomic fragment.

TGN is designed to integrate new data sources as they come available. The ability to do this was tested prior to release by adding genetic association results from a public release of UK Biobank summary statistics for more than 2000 phenotypes, and more recently with the introduction of pQTL data (Yao 2018, Emilsson 2018) into an earlier internal release of the software.

### Computational assist via credible sets

Associations are represented as ‘credible sets’ of variants potentially driving the association (i.e., a set of variants in strong LD that show similar levels of association, any of which could credibly be the functional variant driving the association). Phenotype associations with intersecting credible sets may share the same driver variant - an observation which has particular importance to the development of therapeutic hypotheses in instances where disease associations colocalize with eQTLs, pQTLs or traits that may represent potential new indications or safety risks. Accordingly, while a different index SNP may be listed in different studies due to statistical fluctuation and the widespread LD in the human genome, TGN makes it easy to see whether or not these could represent the same underlying genetic risk factor. For each association or pQTL result, users can select a relevant 1KGp3 population from which to calculate credible sets having r2 >= 0.6 to the index variant.

High density association results from UK Biobank are pre-processed into LD-independent index variants against 1KGp3 EUR reference data, with each retained index variant representing the most significant result (pval <= 1e-6) in its LD neighborhood. Similarly, those high density cis-acting eQTL results from GTEx, which pass tissue-specific significance thresholds, are distilled into LD-independent index variants.

Many publications and user-supplied associations may now come with more detailed fine-mapping information performed based on analysis of raw genotype data (Huang 2017) or complete summary statistics from meta-analyses (Benner 2016). Since such analyses offer a more refined mapping beyond the LD analysis of the index association, these credible sets may be entered by the user, with individual variant posterior probabilities that can be used in all visualizations and analyses instead of the automatically computed index SNP version above.

### Software framework

A main goal in developing TGN is to share it widely so as to encourage collaboration in interpreting and curating targets across academic, consortia and commercial entities. To enable this, the components used needed to be freely available and easy to install. All recovered data are stored in an SQLite database dedicated to the specific target gene. SQLite was chosen for its simple implementation and ease by which databases can be distributed and shared. Data are stored with reference to source and timestamped to permit versioning and to help track data differences across Ensembl releases.

A Java web server based on Spark (sparkjava.com) provides interactive browser-based content driven by Javascript. Through the interface, users can view, annotate, configure, modify, and supplement graphic and tabular representations of collected data for each target gene. The interactive display allows users to rapidly identify association, pQTL, and eQTL results that share credible-set markers. The interface also allows users to add new association, pQTL, and eQTL results, define custom credible sets, and add PubMed, bioRxiv, file, and web-site-based references.

Additionally, multiple graphic excerpts/snapshots (Details) can be stored and annotated with each reference. Details can be directed to appear in different sections of the TGN interface. To enter a detail a user selects/enters the source and can then simply paste information from the clip-board and write a comment; this reduces any barrier to collecting all the information about a target in one place while still connecting it to the source material from which it was captured. Example target databases can be explored at http://tgn.broadinstitute.org/.

## Results

### Enabling efficient data curation for scientists

After the automated database build for a target gene, the data are available for curation. TGN is designed to enable scientists to record and analyze the information as quickly as possible. Scientists with diverse specialties can weigh in with analyses or expert views on different data types – for example, a data analyst may need to evaluate the statistical methods and power regarding a genetic association finding, while a chemist may need to judge whether or not a particular variant will affect the ligand binding in the crystal structure.

In contrast to showing the overwhelming amount of information gathered by the database-build script, TGN initially hides all analysis results. The user can choose to show just the information their expertise judges solid and relevant to the target. However, all other data on the target are still in the interface and other users can see information with curated comments, perhaps marked as “superseded by later meta-analysis” or “sample size only 300”, and know that those references have been reviewed and not missed. TGN also serves as a checklist so users can quickly see which sources have been reviewed and which standard ones are still awaiting review.

### Interface Sections

The TGN interface is divided into a number of sections, with all but the first being partitioned into separate collapsible accordions.

The Genome Display area at the top of Figure 1 shows user-selected data elements aligned to regional genomic landmarks, rendered either in Credible-set mode of LD-summary mode. In Credible-set display mode (shown), each user-displayed association, pQTL, and eQTL result is shown as a labelled set of tick marks, depicting the result’s index variant and any credible-set members. Nonsynonymous coding variants are shown in red. The tick-mark patterns of credible sets quickly enable one to see if two results with different index SNPs likely represent the same underlying genetic cause, evidenced by intersecting credible-set content. Hovering the cursor over any SNP highlights it in turquoise in all results in which it is included. Clicking on the display name for any result will reveal a table below the display area which describes all credible-set members.

**Figure 1:**
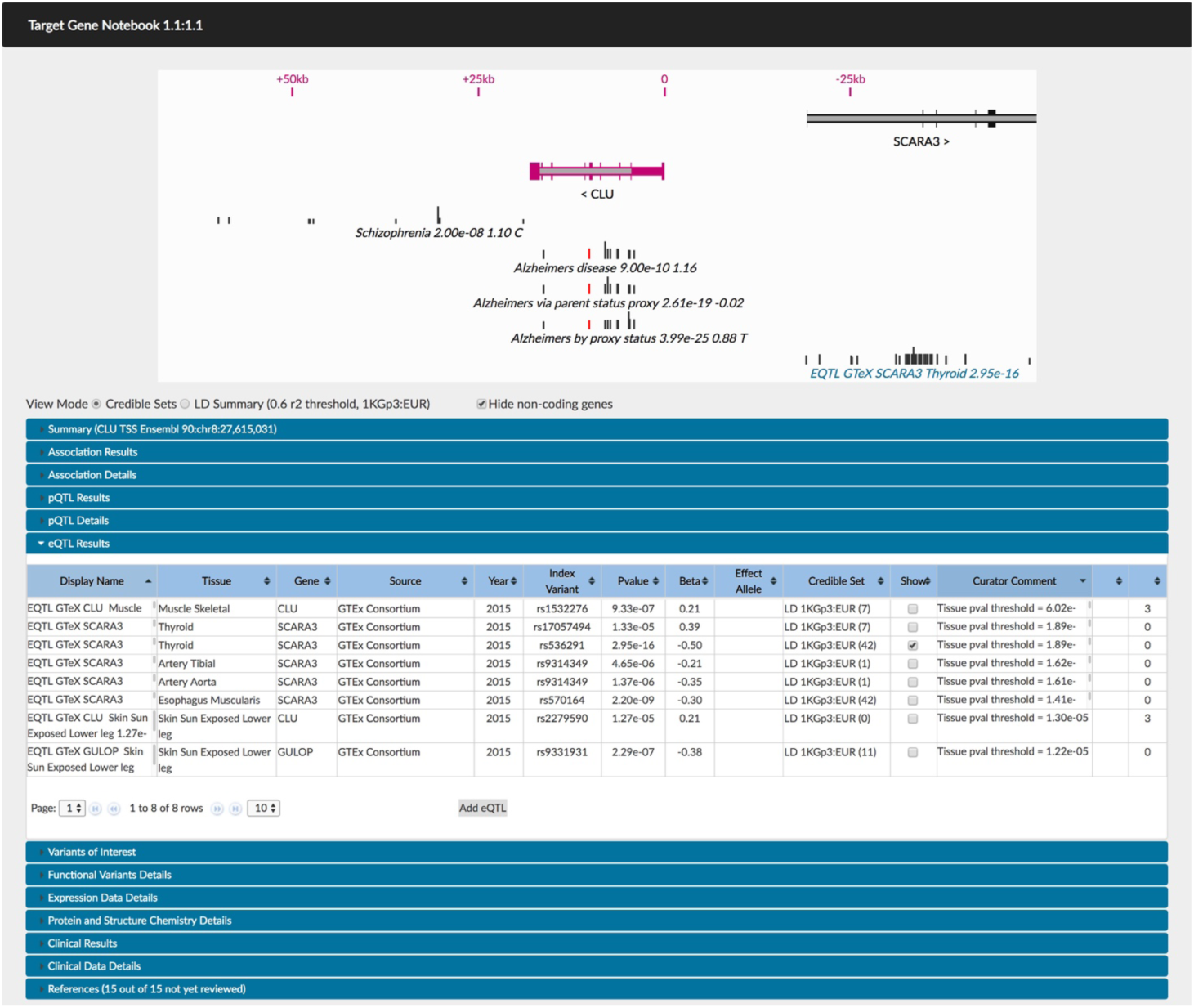
Overall display of data sections for one target (CLU) in Target Gene Notebook. The horizontal tracks of SNPs allow the user to quickly see this is the same association even though different index SNPs are reported in the literature. In this overview all but the eQTL sections are collapsed; other sections can be opened similarly to the eQTL section for display of in-depth information.

The next section, the Summary section, allows users to record general notes and conclusions regarding the target gene and/or local genome region. When others come to look at the database this is the first stop to understand where the in-depth review of the gene is currently at. The Association Results, pQTL Results, and eQTL Results sections below hold a tabular presentation of automatically harvested and user-added GWAS, pQTL, and eQTL results, respectively. Users can select which results are depicted in the Genome Display, edit the text used to label the result in the Genome Display, select or define credible marker sets for each result, and store annotations for each result. Because there can be hundreds of significant eQTL results in a target region, to help their prioritization, the eQTL table has an extra column indicating the number of displayed association and pQTL results possessing intersecting credible sets with the 1KGp3 EUR credible set of each eQTL result.

Furhter down, the Variants of Interest section presents a table of all variant alternate-alleles having a frequency >= 0.001 in at least one of the gnomAD reference populations and also having a coding impact on at least one canonical transcript in the region. Here, “coding impact” is defined as having a Sequence Ontology consequence term amongst these: protein altering variant, missense variant, inframe deletion, inframe insertion, start lost, stop lost, frameshift variant, stop gained, splice acceptor variant, splice donor variant. Each gene impacted by an allele is presented as a separate row in the table. The table also gives users the ability to include or exclude a variant in/from the LD-summary display.

The Clinical Results section contains tables of discovered phenotype-gene and phenotype-allele associations.

The LD-summary display mode (Fig 2) shows genomic landmarks as well as displayed association-, pQTL-, and eQTL-index variants grouped with user-selected common coding-variants such that any variant in a group has an r2 >= 0.6 (against 1KGp3:EUR) to at least one other variant in the same group. Clicking on any LD-grouped set of variants will expose a table of result details for that group below the display area.

**Figure 2:**
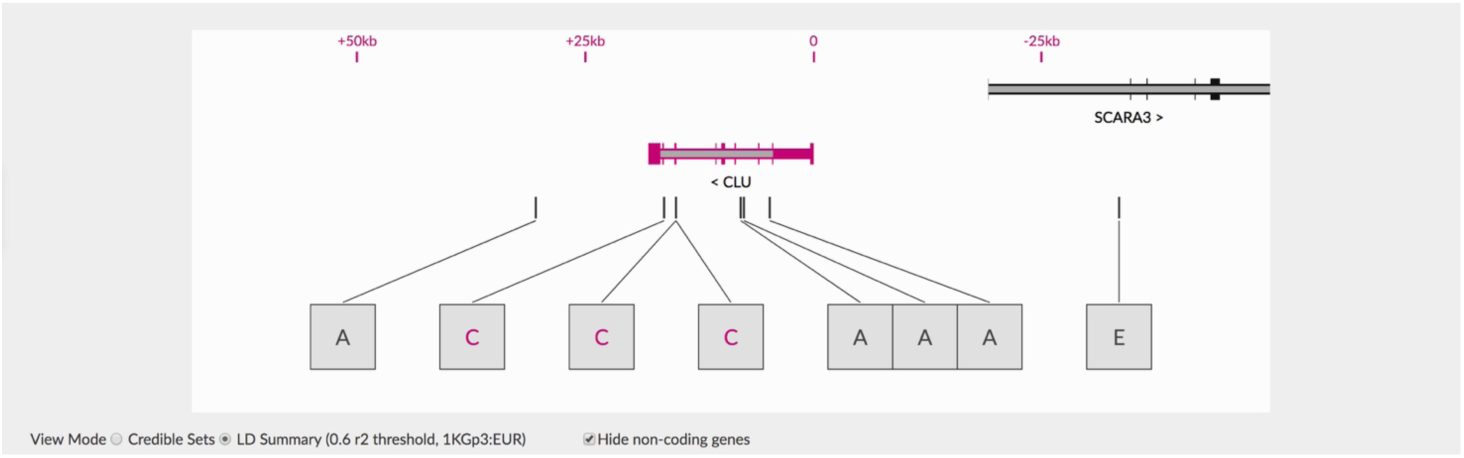
. Alternate view of LD in the CLU region, here showing blocks of LD rather than tracks. A indicates an Association in that block, E an eQTL and P (not shown) a pQTL. Coding variants are indicated by C..

The References section (Figure 3) organizes PubMed, bioRxiv, web-based, and file-based publications related to the target-gene data. It is prepopulated with sources involved in the initial database generation, but users can make additional entries. The status of each reference can be declared with respect to being completely reviewed. Additionally, users can attach images (Details) to each reference, along with descriptions and assignment of the Detail to other interface sections.

**Figure 3.**
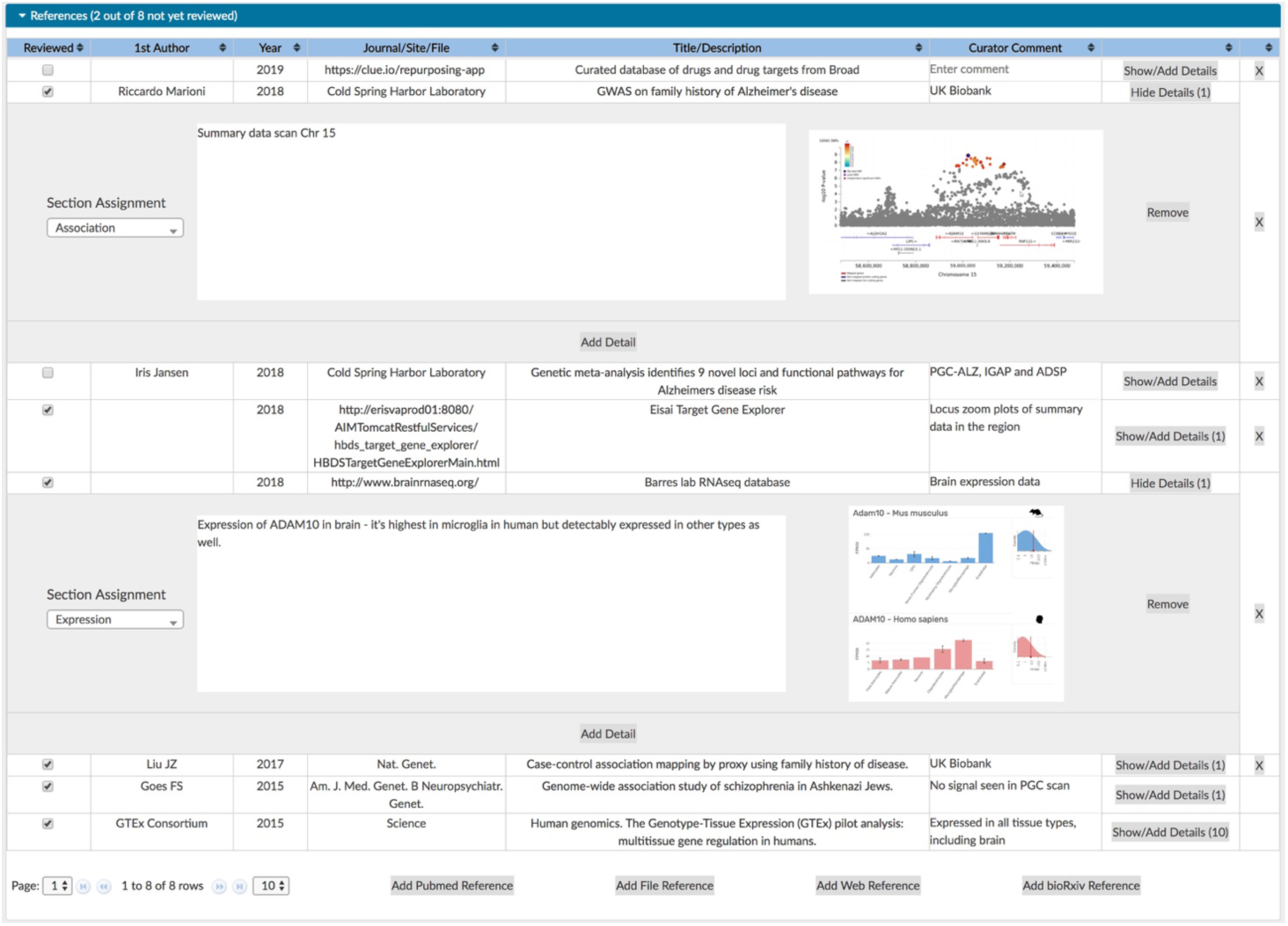
Example of the references and details saved for ADAM10. This section records all the data sources used in the TGN database and provides users a simple method of storing screen captures of related information.

### Database organization

The root web page served by the TGN backend lists all available target-gene databases. Basic location and gene-neighborhood information is shown for each target, as well as the number of database references yet to be declared as fully reviewed. Various keyword and phrase tags can also be assigned to each target database allowing easier filtering and grouping of available databases. The root web page offers a link to a tag-management page where tags can be created and assigned.

## Availability and Future Directions

By distributing TNG as an open-source application our hope is that others may not only benefit from using the application as distributed, but also that it could be a community-based application where new extensions and integrated data types can be added to extend the framework provided. A few other areas of note for further development:

- R2 calculations are performed using the Ensembl API, which, as of Ensembl API v93, calculates LD marker-wise, considering only the reference allele and first alternate allele.
- Data-source harmonization will not always be perfect. For example, allele strandedness is not always able to be fully audited between data sources based on available information.
- User write-events to target-gene databases are not currently pushed to other observers of the same target gene, limiting the user experience when multiple users are simultaneously curating the same target gene.
- User-level permission stratification is not yet implemented, which would allow certain users database-write access, while restricting others to read-only access.
- Certain genomic regions are problematic for TGN, either for LD-calculation aspects (sex chromosomes, HLA) or because of under-curated analysis results (sex chromosomes).

